# Gluconate acts as a signal to induce biofilm in *Bacillus subtilis*

**DOI:** 10.64898/2026.09.08.750294

**Authors:** Courtney E Price, Amelia Sadlon, Amina Bradley, Claudia Perez, Elizabeth A Shank

## Abstract

*Bacillus subtilis* is the best-studied Gram-positive microorganism, but little is known about how the presence and utilization of diverse carbon sources impacts its production of biofilm. Here, we have identified gluconate as a carbon source that induces biofilm wrinkling in *B. subtilis* colony biofilms. Targeted phenotypic analysis revealed that the genes encoding the canonical structural biofilm components (*tasA*, *epsA-O*, and *bslA*) are required for this response. Import and metabolism of gluconate, however, are not required to induce these phenotypic changes. We show that the effect of gluconate on *B. subtilis* is linked to iron availability: production of the siderophore bacillibactin is decreased by gluconate and *B. subtilis* mutants unable to import bacillibactin grow better and wrinkle more in the presence of gluconate. In addition, supplementation of single-carbon media with iron dramatically increased growth and wrinkling of a *B. subtilis* bacillibactin mutant grown on gluconate but did not impact *B. subtilis* grown on glucose. We conclude that gluconate acts as a signal to increase biofilm in *B. subtilis* and increases iron availability in a bacillibactin-independent manner.

**Importance:** *Bacillus subtilis* is a model organism and beneficial bacterium with plant growth-promoting properties. It is also a saprophyte, playing an important ecological role in breakdown of soil organic material. The soil environment contains diverse carbon sources and metabolites that act as signals to regulate bacterial behavior, but the complexity of this environment limits what is known about how individual factors shape microbial communities. We have used a screening approach to identify a novel carbon source, gluconate, that regulates *B. subtilis* biofilm formation and supports iron acquisition. Gluconate is a byproduct of glucose metabolism by bacteria and fungi, thus indicating a role for gluconate in mediating *Bacillus* community interactions. Overall, these results demonstrate new insights into how the complex soil environment shapes bacterial behavior.

## Introduction

*Bacillus subtilis* is a well-characterized bacterium used extensively in both basic research and biotechnology settings (1–3). *B. subtilis* is an important model for understanding biofilm development: the process by which microorganisms transition from a planktonic to a sessile lifestyle during which they grow as a heterogeneous community encased in a self-produced extracellular matrix (4–6). This sessile biofilm lifestyle protects microbes from environmental challenges such as oxidative stress, predation, and antibiotics, and is the predominant bacterial lifestyle in the environment. Although domesticated laboratory strains lack many group behaviors, the study of emergent *B. subtilis* behaviors, including biofilm formation, is facilitated by the use of wild, undomesticated *B. subtilis* strains (7–10). The undomesticated strain that we use throughout this study, *B. subtilis* NCIB3610, is ancestral to the widely used, genetically competent laboratory strain *B. subtilis* 168 and is commonly used to study biofilm formation (7, 11–13). *B. subtilis* biofilms are composed of three major structural components: the fiber-forming protein TasA, an exopolysaccharide (EPS), and a hydrophobin (BslA) that imparts hydrophobicity to the biofilm (4–6, 14–16). *B. subtilis* biofilms are commonly studied in two forms – either as a floating pellicle formed at the air-liquid interface, or on an agar surface as a colony biofilm (4, 17, 18).

Regulation of *B. subtilis* biofilm formation is a complex process coordinated by the master regulator Spo0A, a transcription factor that serves as an integration point for several upstream response regulators and controls the transition into biofilm gene expression and sporulation (4–6, 19). Biofilm gene expression is largely dependent on the levels of phosphorylated Spo0A (Spo0A∼P) within the cell (20–22). The sensor histidine kinases KinA, KinB, KinC, and KinD are major contributors to the phosphorylation state of Spo0A and are responsive to diverse environmental signals (23). There is substantial redundancy among these four kinases, and phenotypic responses are condition dependent (24, 25).

Metabolism has a critical role in shaping *B. subtilis* biofilm development, and environmentally available carbon has been shown to impact *B. subtilis* biofilm formation through diverse mechanisms (26–33). Biofilm formation can be modulated by intermediaries generated during carbon catabolism, some of which feed directly into the production of biofilm components, while other intermediate catabolism products can build up to levels that inhibit and disrupt normal metabolism (26–28). Alternatively, some carbon sources, such as those associated with plant roots, can act as environmental signals that alter biofilm matrix synthesis (29, 30, 34). Several plant root polysaccharides induce biofilm matrix production in a KinCD-dependent manner (29). Similarly, KinD mediates the signaling response to glycerol and manganese, two components in the commonly used biofilm-inducing medium MSgg (9, 30). Carbon source signaling is not exclusively regulated by the sensor kinases, as xylan also induces biofilm via an unidentified, Kin-independent mechanism (29), while lactate regulates biofilm formation through AI-2 quorum sensing (34). Taken together, these studies demonstrate a strong but not-well-understood link between carbon source metabolism, signaling, and biofilm formation in *B. subtilis* that appears highly specific to the carbon source examined.

Biofilm formation is also closely linked to iron acquisition in *B. subtilis*, as iron acquisition is required for robust biofilm formation. Bacterial iron uptake is mediated by siderophores, small self-produced iron chelators that bind iron and then are taken back up by the bacteria (35, 36). In *B. subtilis*, biofilm matrix production facilitates efficient uptake of iron-bound siderophores, and matrix-associated iron acts as an extracellular electron acceptor during respiration (37–39). *B. subtilis* encodes one siderophore, bacillibactin, as well as several pathways for the uptake of other forms of iron or siderophores pirated from other microorganisms (35, 40, 41). Although iron scavenging has been linked to metabolic changes due to co-regulation of the iron-sparing response and metabolism (42), and iron starvation also triggers the stringent response (43), there is little understanding of how specific carbon sources alter iron acquisition in *B. subtilis* and what interactions may exist between carbon source, iron acquisition, and biofilm formation.

Previous studies examining the biofilm response of *B. subtilis* to carbon source have focused on carbon sources associated with plants (*B. subtilis*’ presumed natural environment) or carbon sources commonly used in laboratory settings (i.e., glucose). Here, we took an untargeted approach to identify carbon sources that induce biofilm in *B. subtilis* without making *a priori* assumptions about their relevance to *B. subtilis*. This allowed us to identify gluconate, a simple sugar that is produced by soil-resident bacteria and fungi (44–47), as a strong inducer of *tapA* gene expression (*tapA* is the first gene in the TasA operon encoding the fiber-forming biofilm component) in liquid culture as well as phenotypic wrinkling in agar colony biofilms. We show that gluconate metabolism is not required for biofilm induction, nor is this response dependent on the sensor histidine kinases (KinABCD). Instead, we provide evidence that gluconate increases iron accessibility to the cell without itself being imported: *B. subtilis* grows better in the absence of bacillibactin import when cells are grown on gluconate. We have therefore identified a novel link between biofilm formation, carbon utilization, and iron access in *B. subtilis*.

## Results

### Expression of the biofilm gene *tapA* is induced by a specific, limited set of carbon sources

Previous research has shown that carbon sources from plant root exudates induce *B. subtilis* biofilm formation (29). To gain a better understanding of how diverse carbon source utilization is linked to biofilm gene expression in *B. subtilis*, we grew a luminescent *tapA* transcriptional reporter strain, a proxy for biofilm formation, with 190 individual carbon sources in a carbon-source-free base minimal salts nitrogen (MSN) medium. We monitored bacterial growth and *tapA* expression over time to identify novel carbon sources that increased *tapA* gene expression (Table S1; Figure S1). We identified 17 candidate carbon sources that induced *tapA* expression in the initial screen. We re-tested each carbon source at 0.5% w/v alone and with 0.5% D-glucose (glucose) or 0.5% D-cellobiose (cellobiose) (Figure S2). We confirmed that *B. subtilis* had significantly higher *tapA* expression when grown in five of the carbon sources identified in the initial screen relative to growth in glucose or cellobiose alone (Figure 1A; Table S2): D-gluconic acid (gluconate), D-galacturonic acid, L-ornithine, L-proline, and L-histidine. Additionally, *tapA* expression was significantly higher in L-aspartic acid plus cellobiose than in cellobiose alone (Figure S2, Table S2). D-galacturonic acid was previously reported to induce biofilm gene expression in *B. subtilis* (29), but gluconate has not been shown to impact *B. subtilis* biofilm formation and was the strongest inducer of *tapA* expression identified in this screen. Interestingly, gluconate induced *tapA* more strongly as a sole carbon source than in the presence of cellobiose or glucose (Figure 1). In contrast, D-galacturonic acid induced *tapA* most strongly in the presence of cellobiose but did not strongly induce *tapA* as the sole carbon source (Figure 1).

**Figure 1.**
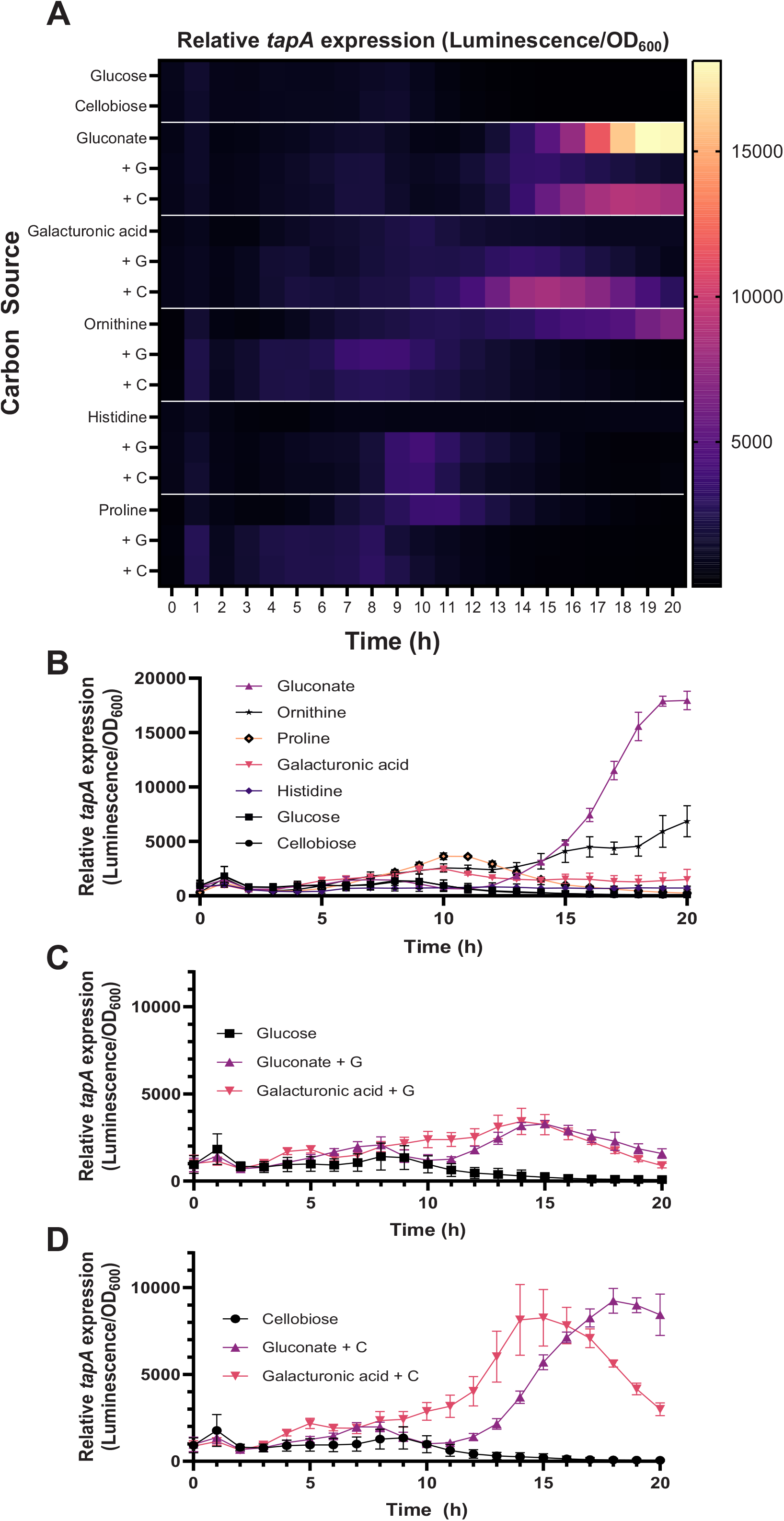
Expression of *tapA* is induced by several candidate carbon sources. *B. subtilis* 3610 *sacA*::P*tapA*-LUXABDCE was cultured in liquid MSN in the presence of each candidate carbon source at 0.5% w/v. *B. subtilis* was also cultured with each candidate carbon source plus 0.5% w/v glucose (+G) or 0.5% cellobiose (+C). Growth (OD600) and Luminescence were measured hourly over a 20-hour time course. Luminescence was normalized to growth at each time point. **A)** Each box represents the mean of 4 biological replicates. **B-D)** Each point represents the mean and standard deviation of 4 biological replicates for assays in **B)** single candidate carbon sources, and candidate carbon sources with **C)** glucose or **D)** cellobiose. Statistical results are available in Table S2.

Because biofilm and sporulation are both controlled by the central regulator Spo0A in *B. subtilis* (20–22), we also tested how gluconate impacted the production of *B. subtilis* spores (Figure S3). The percentage of spores was significantly lower in gluconate relative to glucose or cellobiose at 24 hours, not significantly different between carbon source conditions at 48 hours, and significantly higher in gluconate by 72 hours (Figure S3). The presence of two carbon sources led to a significantly lower percentage of spores in the population at all time points. We also observed the formation of a brown pigment in the gluconate culture at 48 and 72 hours (Figure S3), which is consistent with the sporulation-associated pigment pulcherrimin (48–52). These results indicate that both *B. subtilis* biofilm gene expression and sporulation can be markedly induced by gluconate and that the presence of multiple carbon sources plays a role in modulating the response of *B. subtilis* to individual carbon sources.

### Gluconate induces phenotypic wrinkling in *B. subtilis* colony biofilms

Because gluconate had the strongest impact on *tapA* expression and had not been previously identified to impact *B. subtilis* biofilms, we next tested whether gluconate as a sole carbon source induced a phenotypic response in *B. subtilis* colony biofilms. To do so, we grew *B. subtilis* colony biofilms on MSN agar containing 0.5% glucose, 0.5% cellobiose, or 0.5% gluconate and imaged after 24, 48, and 72 hours of growth (Figure 2A; Figure S4). The diameter of *B. subtilis* colonies grown on glucose was significantly smaller than the diameter of colonies grown on gluconate or cellobiose (Figure S4A-C). This size difference was not due to differences in growth, however, as the number of colony forming units (CFUs) were not significantly different at any timepoint between glucose and gluconate (Fig S4D). Colony morphologies were distinct between carbon sources at each time point (Figure 2A). By 48 hours of growth, colonies on gluconate displayed a wrinkled phenotype in the center of the colony indicative of biofilm formation, which was also apparent at 72 hours. To determine whether an additional, second carbon source would alter the wrinkling phenotype induced by gluconate, we examined colony biofilm morphology on MSN with gluconate plus either glucose or cellobiose (Figure S5). Gluconate plus cellobiose led to less phenotypic wrinkling, while gluconate plus glucose led to minimal wrinkling. This is consistent with the results from liquid culture, where *tapA* expression was more modestly induced by gluconate plus cellobiose and was not significantly induced by gluconate plus glucose (Figure 1A).

**Figure 2.**
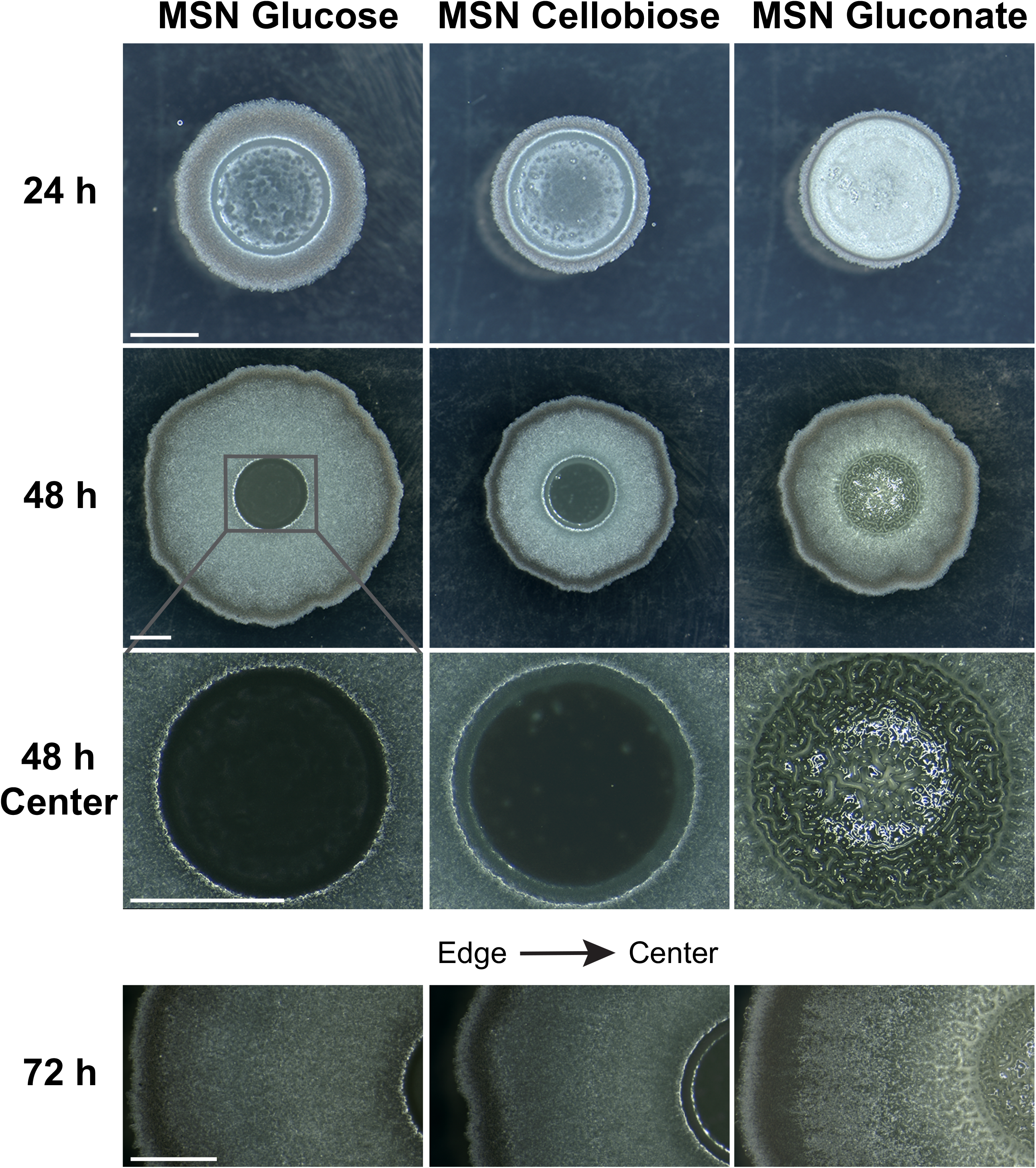
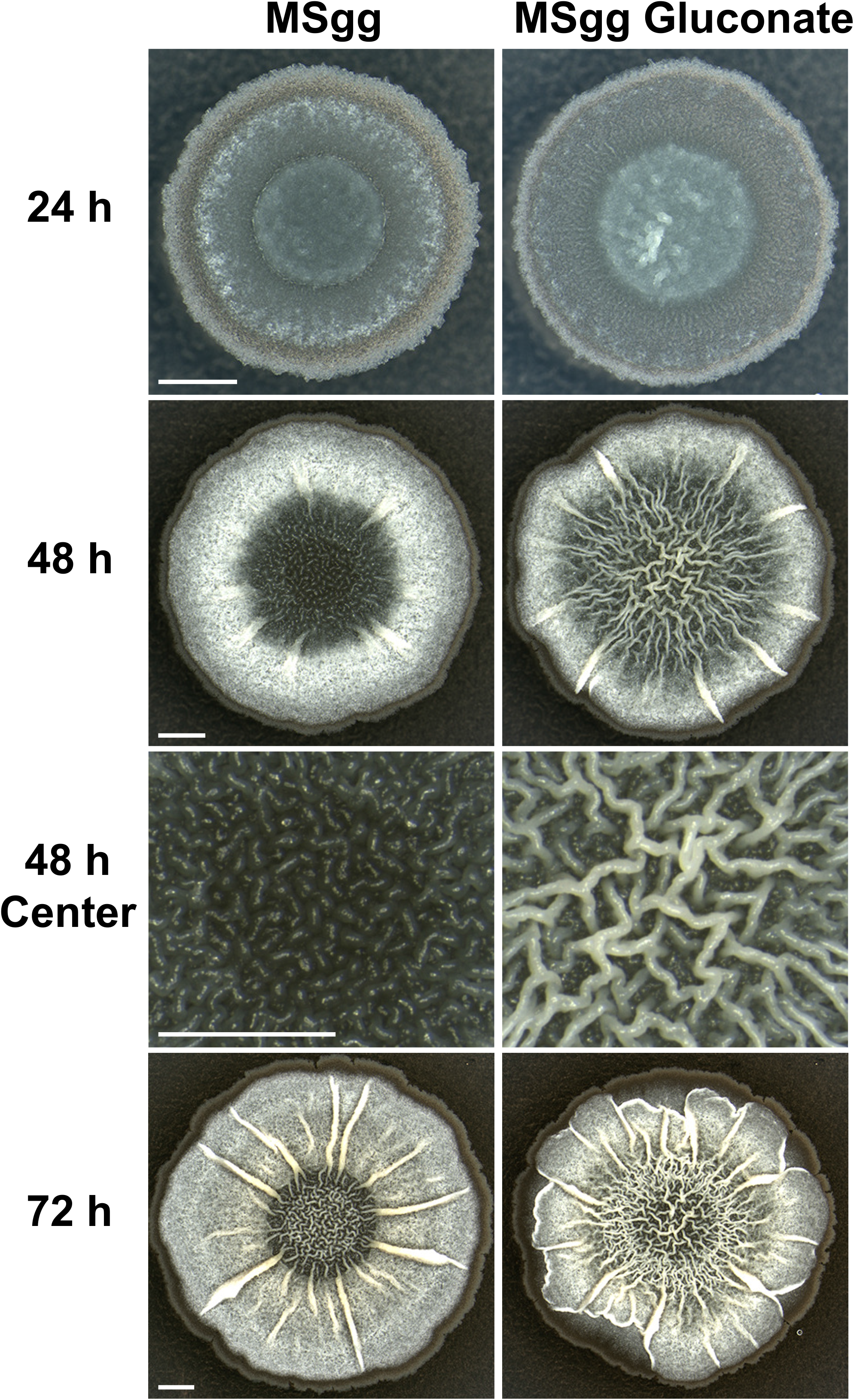
Gluconate induces phenotypic wrinkling in *B. subtilis*. Representative brightfield images (n=3-6 biological replicates) of *B. subtilis* 3610 colony biofilms after 24, 48, and 72 hours of incubation on **A)** MSN with 0.5% w/v glucose, cellobiose, or gluconate and **B)** MSgg with or without 0.5% w/v gluconate. Scale bars indicate 2 mm, and the scale is consistent across each row within the same base medium.

We next tested whether the addition of gluconate to a common biofilm-inducing medium, MSgg, which contains glycerol and glutamate as carbon sources, also increased wrinkling. The addition of gluconate to MSgg caused dramatic changes in *B. subtilis* colony biofilm morphology, with more and smaller wrinkles visible at 24, 48, and 72 hours of growth (Figure 2B). These results demonstrate that gluconate can induce phenotypic wrinkling under multiple media conditions, whether as a sole carbon source or in the presence of other carbon sources.

### Gluconate-induced wrinkling is dependent on *B. subtilis* biofilm genes

Phenotypic wrinkling in *B. subtilis* colonies is typically due to biofilm formation, so we next tested whether the wrinkling induced by gluconate was dependent on canonical biofilm genes. We first examined the morphology of the matrix-deficient mutant *B. subtilis epsA-O*::*tet^R^ tasA*::*kan^R^ bslA*::*erm^R^,* which cannot produce any of the three major *B. subtilis* structural biofilm components, on both MSN and MSgg agar supplemented with 0.5% gluconate (Figure 3). This mutant did not wrinkle under any condition, indicating that these matrix genes are required for the phenotypic response to gluconate.

**Figure 3.**
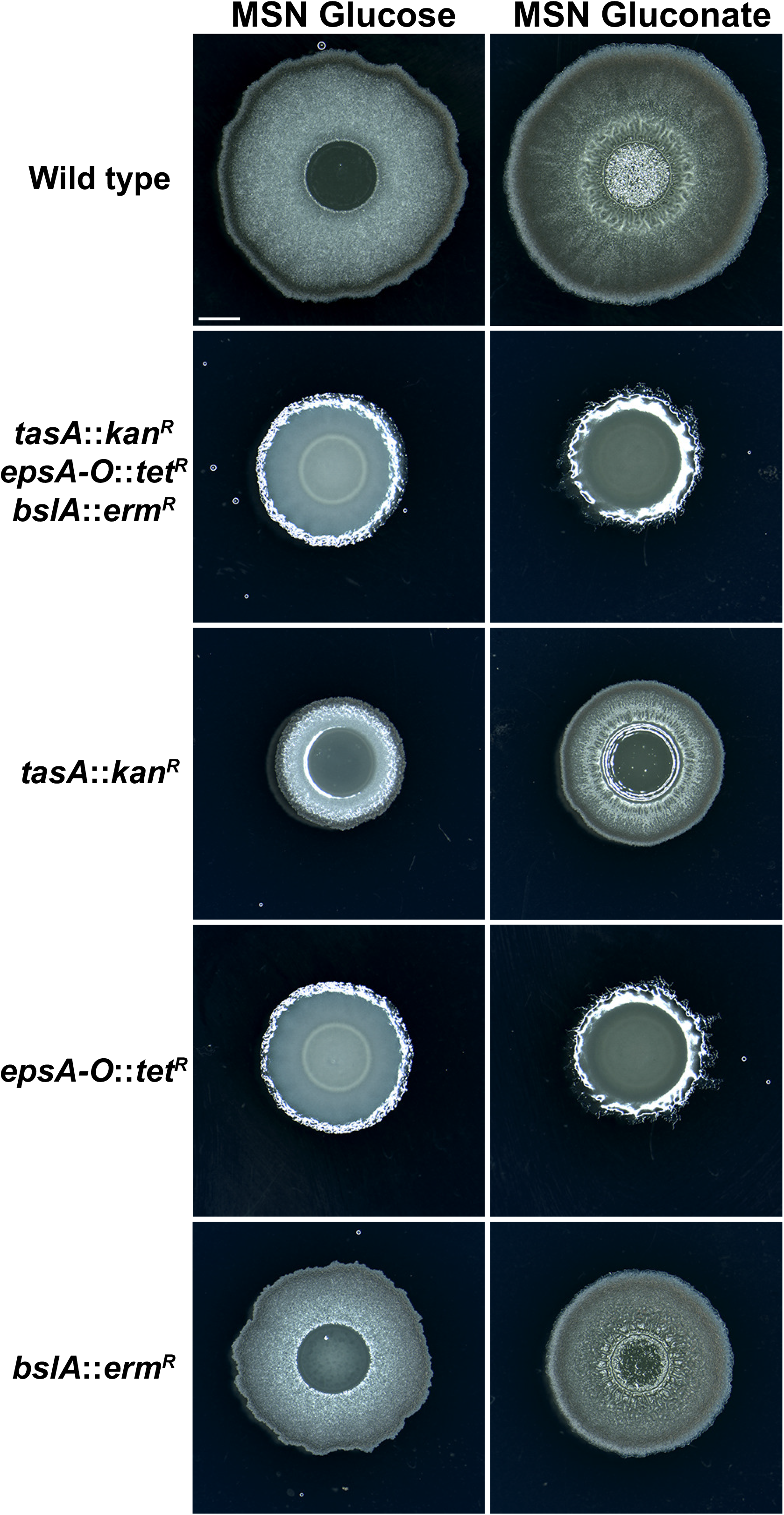
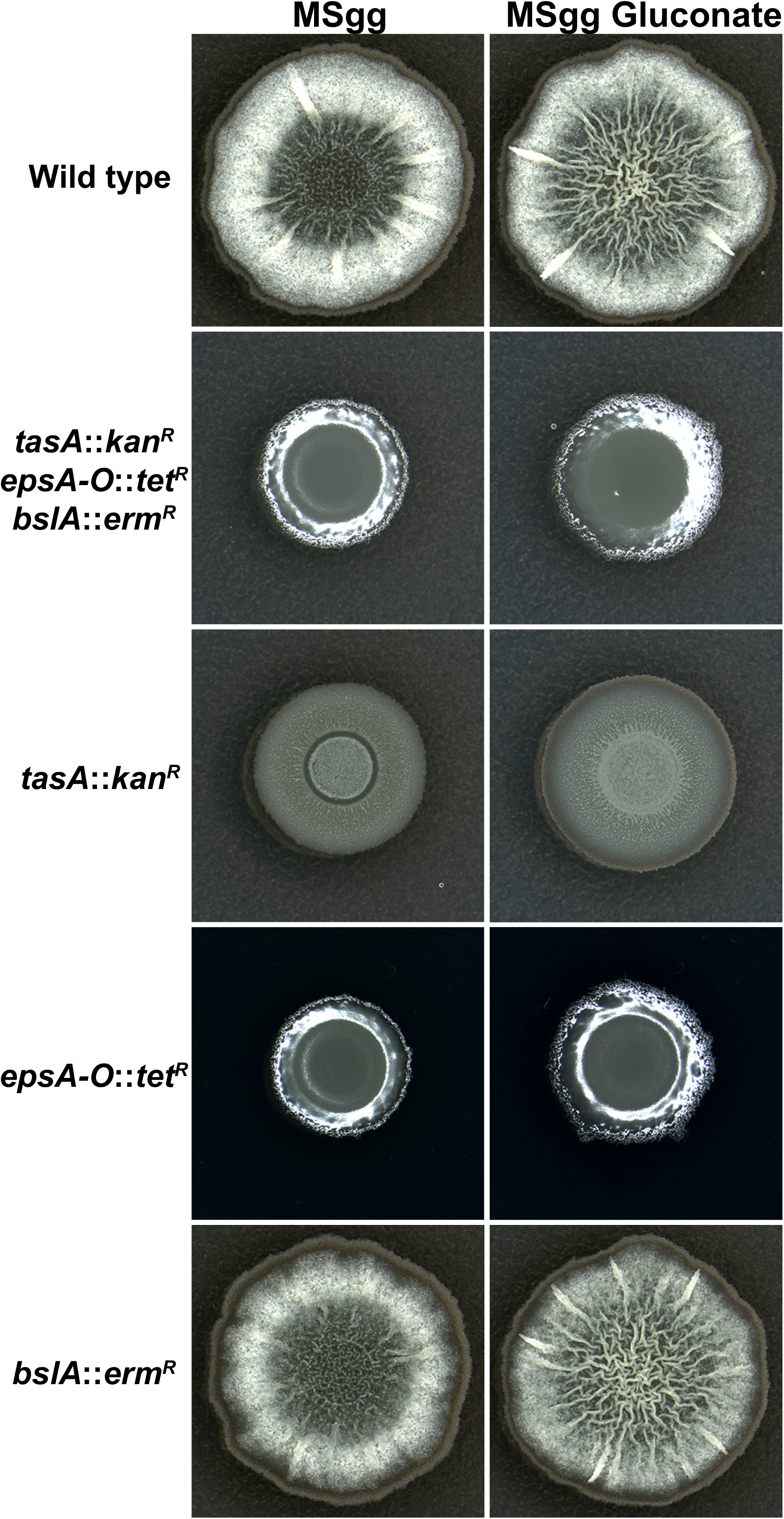
Biofilm genes are required for *B. subtilis* to wrinkle in response to gluconate. Representative brightfield images (n=3-6 biological replicates) of *B. subtilis* 3610 colony biofilms after 24, 48, and 72 hours of incubation on **A)** MSN with 0.5% w/v glucose, cellobiose, or gluconate and **B)** MSgg with or without 0.5% w/v gluconate. Deletion strain genotypes are indicated on the left, and all gene deletions are replaced with one of the following resistance cassettes: kanamycin (kan^R^), erythromycin (erm^R^), or tetracycline (tet^R^). The scale bar indicates 2 mm and is consistent across all images.

To determine whether a specific biofilm component or multiple components were required for wrinkling in response to gluconate, we also tested *B. subtilis* mutants missing single genes for each individual matrix component: *tasA*, *epsC*, and *bslA*. All three biofilm components were required for phenotypic wrinkling in response to gluconate on MSN, where gluconate was the sole carbon source and MSN is not known to induce biofilm formation. However, on MSgg, where there are two other carbon sources and the medium induces biofilm formation independently from gluconate, *tasA* and *epsC* were required for the response to gluconate, but *blsA* was not (Figure 3). Strains deleted for other genes within the biofilm operons, including *tapA, sipW,* and *epsA,* also did not wrinkle on MSN gluconate (Figure S6). Furthermore, deletion mutants of major regulators of biofilm formation, including *spo0A, sigH,* and *remA*, similarly did not wrinkle on MSN gluconate (Figure S6). These results indicate that the morphological wrinkling in *B. subtilis* colonies induced by gluconate is dependent on the presence and expression of the canonical *tasA* and *eps* biofilm genes through characterized regulatory pathways.

### Gluconate does not increase the expression of canonical biofilm genes

We next tested whether *tapA* gene expression increased on gluconate agar, as it did in liquid culture. We grew a *tapA* fluorescent transcriptional reporter strain on MSN or MSgg agar with or without gluconate and captured colony images at 18, 24, and 48 hours (Figure 4; Figure S7). Surprisingly, average colony fluorescence of the *tapA* reporter was significantly lower in the presence of gluconate at all time points in both MSgg and MSN.

**Figure 4.**
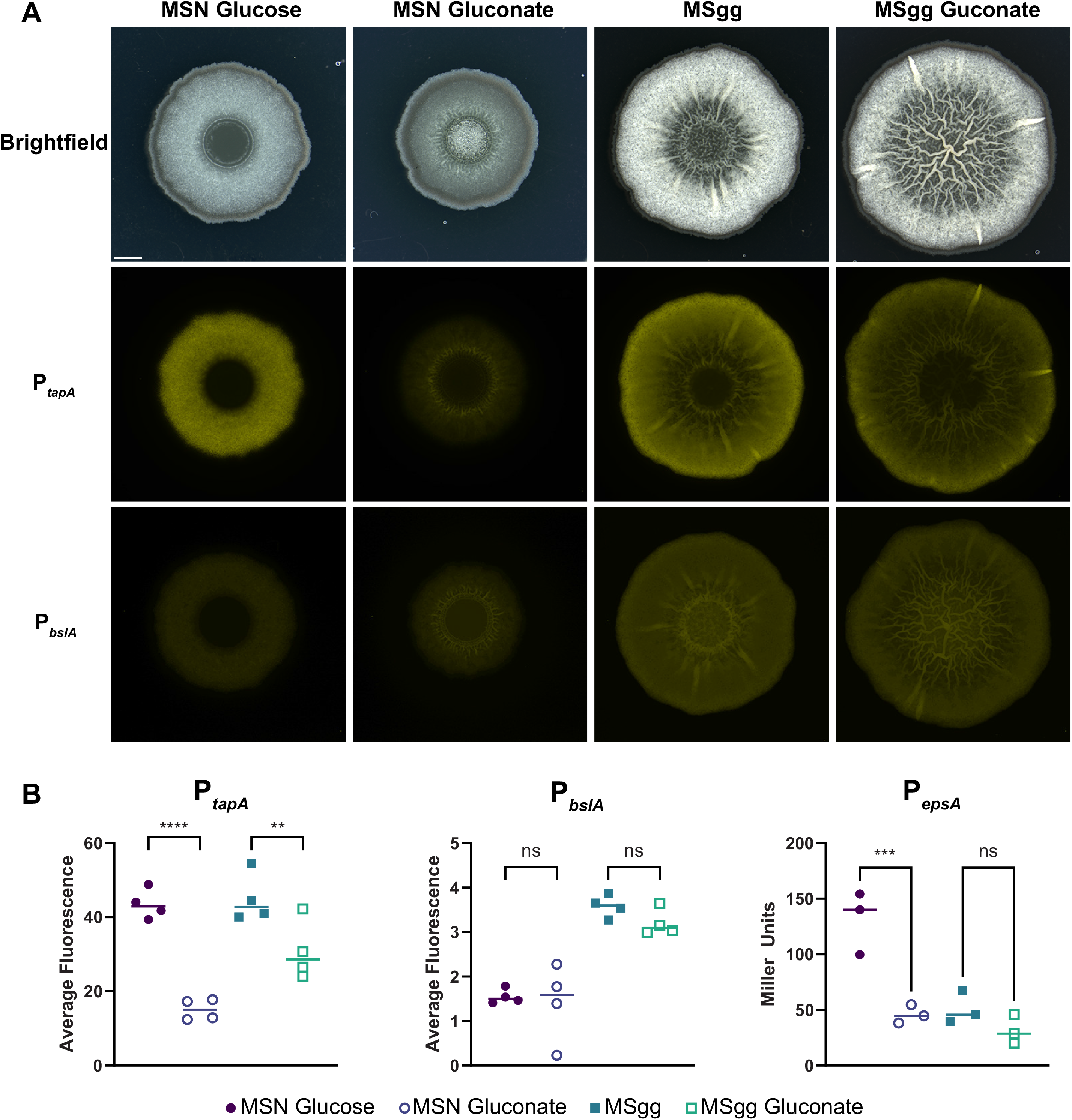
Gluconate does not increase *B. subtilis*’ biofilm gene expression. **A)** Representative brightfield and fluorescence images of *B. subtilis* 3610 colony biofilms after 48 hours growth on the indicated media. Image brightness and contrast adjustments are consistent across all media conditions (rows) but not between reporter strains (columns). **B)** Fluorescence or ONPG (*lacZ*, Miller Units) quantification after 48 hours growth of the indicated reporter strain. Each point represents a single biological replicate. Statistical significance was determined by one-way ANOVA followed by Sidak’s multiple comparisons testing. ** P < 0.01, *** P < 0.001, **** P < 0.0001, ns = not significant.

We next tested whether *B. subtilis*’ phenotypic response to gluconate on agar could be due to expression changes in the two other major biofilm components; EPS and BslA. We constructed a *B. subtilis* fluorescent reporter strain for *bslA* and quantified its fluorescence in colony biofilms (Figure 4; Figure S7). We detected no significant differences in *bslA* expression in most conditions, except at 24 hours in MSgg, where the addition of gluconate led to lower *bslA* expression. To account for the transient and generally weaker expression of the catalytic *eps* genes, we used a *B. subtilis* P*_epsA_*-*lacZ* reporter and quantified *eps* gene expression by ONPG assay. We detected lower or no difference in β-galactosidase activity when gluconate was included in the media, in a timepoint and media dependent manner (Figure S7).

We next wondered whether any changes in biofilm gene expression might be masked due to the many cell layers of thick *B. subtilis* biofilms obscuring fluorescence detectability in top-down fluorescence microscopy images. We therefore quantified the percentage of *tapA-* and *bslA-*expressing cells in colony biofilms grown on MSN and MSgg with or without gluconate using flow cytometry. Similar to our microscopy results, the proportion of fluorescent cells was the same or lower in the gluconate conditions at every time point for both *tapA* and *bslA* (Figure S7). These results confirmed that the number of cells expressing biofilm genes are lower or unchanged when *B. subtilis* is grown as a colony biofilm on gluconate-containing agar, even though gluconate increases wrinkling in a manner that requires the three major *B. subtilis* structural biofilm components.

### Gluconate catabolism is not required to induce *B. subtilis* biofilm formation

We next considered whether the mechanism by which biofilm wrinkling was induced required metabolism of gluconate. *B. subtilis* contains an operon for gluconate utilization that consists of three genes: *gntP*, *gntK*, and *gntZ*. The key functions of these genes are gluconate import (*gntP*), conversion of gluconate to 6-phospho-D-gluconate (*gntK*) and conversion to D-ribulose-5-phosphate + NADPH (*gntZ*) (Figure 5A). We first grew single deletion mutants of each *gnt* gene on MSN supplemented with either cellobiose or gluconate to determine whether each *gnt* gene was required for catabolism (Figure 5B). The morphology of the *B. subtilis gnt* mutants on cellobiose was similar to wild type. Colony size and total cell populations of all of the *gnt* mutants were substantially reduced when grown on MSN gluconate as a sole carbon source (Figure 5B, Figure S8). Interestingly, the *gntP* (gluconate importer) and *gntK* (glucokinase) mutants displayed no wrinkling and almost no growth under these conditions, indicating that gluconate was not metabolized, while the *gntZ* mutant (gluconate dehydrogenase) had slightly more growth and dense wrinkling on gluconate. This increased growth of the *gntZ* deletion strain relative to the *gntP* and *gntK* strains is likely because *B. subtilis* has two other proteins, GndA and YqeC, that are homologs of GntZ. Indeed, GndA appears to be more important for catalyzing the transformation of 6-phospho-D-gluconate than GntZ (53). We next grew the *gnt* mutants on MSN containing both gluconate and cellobiose to determine whether *B. subtilis* would wrinkle in response to gluconate when growth was facilitated by cellobiose, but gluconate was not metabolized (Figure S9). We saw that all three mutants wrinkled in the presence of cellobiose and gluconate. The *gntZ* deletion strain still exhibited the most wrinkling and growth of the three *gnt* mutants and looked more like wild type at both carbon concentrations (Figure S9). We also noted that wild-type *B. subtilis* wrinkled less at a higher total carbon concentration of 1%.

**Figure 5.**
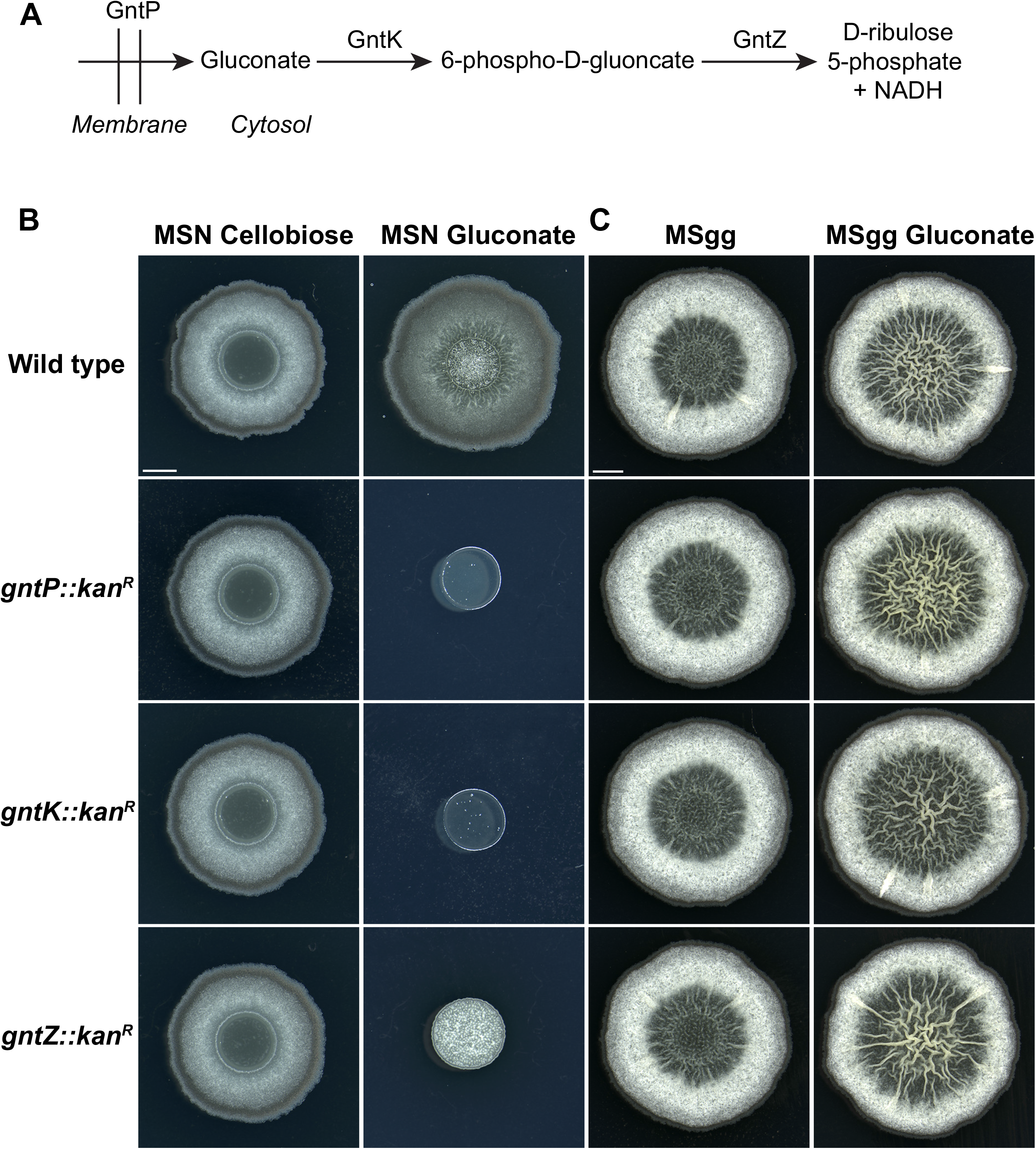
Gluconate acts as a signal to induce wrinkling in *B. subtilis* colony biofilms. **A)** Diagram of the function of *gnt* genes in the import and catabolism of gluconate. **B-C)** Representative brightfield images (n=3-6 biological replicates) of *B. subtilis* 3610 colony biofilms after 24, 48, and 72 hours of incubation on **B)** MSN with 0.5% w/v cellobiose or gluconate and **C)** MSgg with or without 0.5% w/v gluconate. Strain genotypes are indicated on the left, and all gene deletions are replaced with kanamycin (kan^R^). The scale bar indicates 2 mm and is consistent across all images with the same base medium.

Finally, we tested whether metabolism of gluconate was required for the wrinkling response to gluconate on MSgg (Figure 5C). The morphology of the *gnt* mutants on MSgg agar was indistinguishable from wild-type *B. subtilis,* both with and without gluconate. Taken together, these data strongly indicate that gluconate does not need to be imported or catabolized to change colony biofilm morphology and is therefore acting as a signal to induce biofilm wrinkling.

### Deletion of sensor histidine kinases does not impact the response of *B. subtilis* to gluconate

To identify the mechanism underlying the biofilm wrinkling response to gluconate, we examined the morphology of *B. subtilis* mutants lacking key genes involved in the initiation of biofilm formation. We analyzed single and double deletions of four sensor histidine kinases (*kinA*, *kinB*, *kinC*, and *kinD*) to determine whether the response to gluconate depends on these well-characterized regulators. Deletion of these kinases minimally altered *B. subtilis* colony morphology on MSN plus glucose and, unsurprisingly, did not induce wrinkling (Figure S10). Strains with *kinA, kinB* or both deleted wrinkled similarly to a wild-type strain on MSN plus gluconate (Figure S10), indicating that gluconate induces biofilm independently from these two genes. Deletion of *kinC,* but not *kinD*, led to a small increase in wrinkling in response to gluconate, while the double *kinC kinD* deletion strain formed a smaller colony than either single gene deletion, but remained wrinkled (Figure S10). These data indicate that *kinC* and *kinD* are required for normal growth on gluconate and that *kinC* may act as a mild negative regulator of the biofilm response to gluconate. However, none of the four kinases were required for *B. subtilis* to wrinkle in response to gluconate, indicating that they are not required to transduce a positive signal for gluconate-induced biofilm wrinkling, as has been reported for other carbon sources (29).

### Deletion of the Rap-Phr regulators does not impact the response of *B. subtilis* to gluconate

Because we did not identify a positive regulator of the response to gluconate among the sensor histidine kinases, we expanded our candidate approach by examining the morphology of *B. subtilis* strains carrying deletions of the *rap* and *phr* genes. The Rap-Phr phosphatase regulators in *B. subtilis* have been implicated in environmental adaptation, biofilm formation, and heterogeneity in *B. subtilis* (54, 55). Similar to experiments above, we examined the morphology of single-gene deletions of *rap* and *phr* genes on MSN plus gluconate (Figure S11). We did not see any differences in wrinkling relative to wild-type, except for the *phrI* and *phrK* deletion strains (Figure S11, Figure S12), which wrinkled less than wild type. However, both deletion mutants also wrinkled less on MSgg without gluconate (Figure S12). These results indicate that the absence of *phrI* or *phrK* reduces wrinkling on multiple media conditions, but that this reduction is not specific to gluconate. Therefore, the Rap-Phr systems do not appear to transduce a specific positive signal to induce biofilm wrinkling in *B. subtilis* in response to the presence of gluconate.

### *B. subtilis* decreases bacillibactin production in the presence of gluconate

Specialized metabolites drive biofilm formation and differentiation in *B. subtilis* (56, 57), and an increase in specific specialized metabolites is associated with biofilm formation (58, 59). Because our results pointed to an increase in biofilm wrinkling caused by gluconate, but not a clear mechanism by which biofilm was increased, we next quantified specialized metabolites from *B. subtilis* grown on media with and without gluconate by tandem liquid chromatography-mass spectrometry (LC-MS/MS) to determine whether gluconate altered patterns of specialized metabolite production (Figure 6). Five of the seven specialized metabolites that were detected with MSN as a base medium were significantly higher on gluconate than on glucose (pulcherrimin, bacillaene, plipastatin, sporulation killing factor (SKF), and surfactin), consistent with increased metabolite production under biofilm-inducing conditions seen in previous studies (Figure 6A-E) (56, 60). The increase in pulcherrimin, which is associated with the timing of sporulation (48), is also consistent with the increase in sporulation and sporulation pigment that we saw in liquid MSN with gluconate (Figure S3). Two metabolites, bacillibactin and subtilosin, were significantly decreased on MSN gluconate (Figure 6F-G).

**Figure 6.**
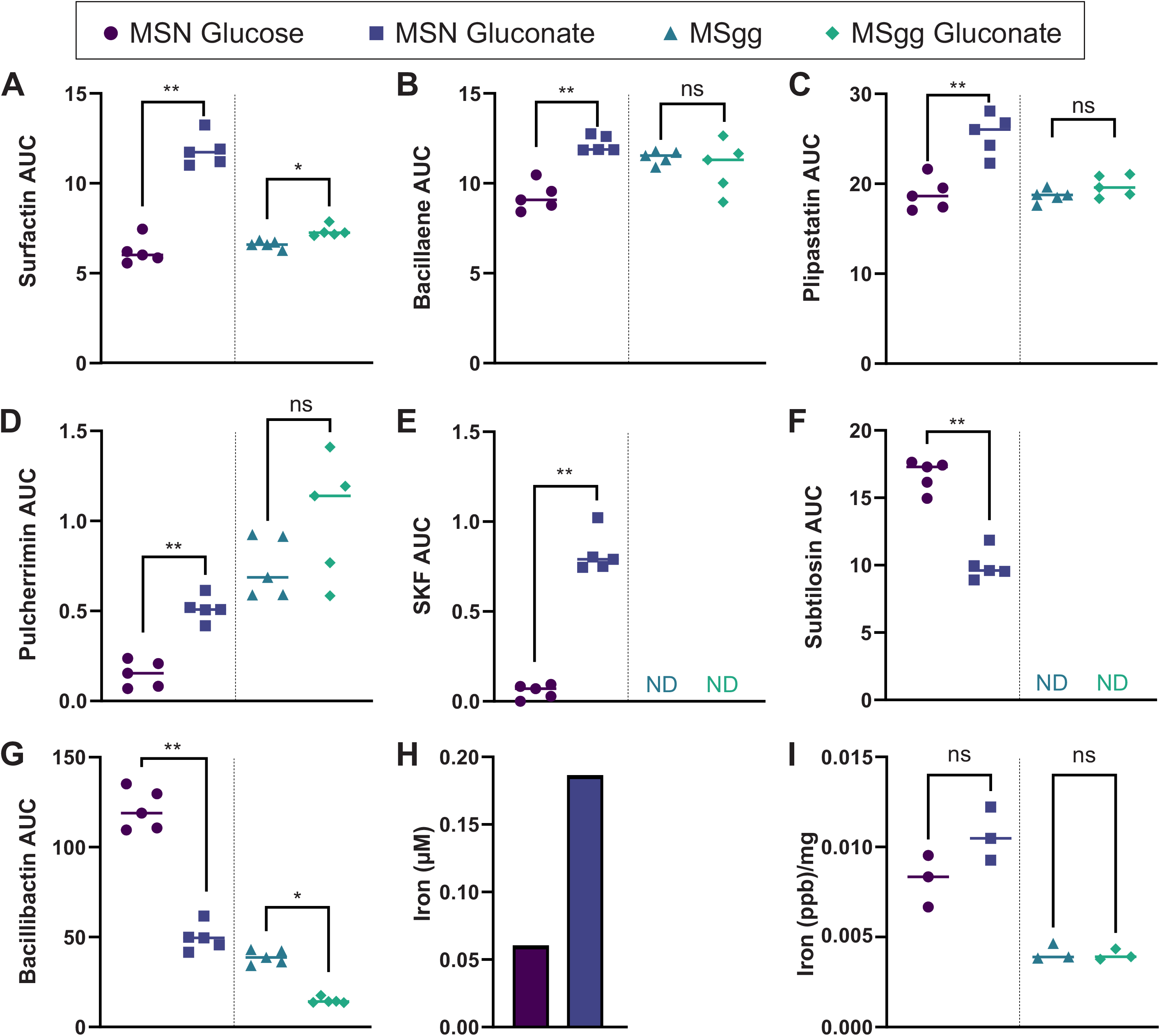
Iron acquisition by *B. subtilis* is altered by the presence of gluconate. **A-G)** LC-MS/MS quantification of metabolites produced by wild-type *B. subtilis* 3610 colony biofilms incubated on MSN with 0.5% w/v glucose or gluconate and MSgg with or without 0.5% w/v gluconate. AUC = Area Under the Curve. Mann-Whitney U tests were performed for each metabolite between paired base media conditions, and a Benjamini-Hochberg multiple comparisons correction was performed. **H-I)** ICP-MS quantification of iron in parts per billion (ppb). **H)** Iron quantification in MSN medium with glucose or gluconate. **I)** Iron quantification in *B. subtilis* colony biofilms after 48 hours of incubation on the indicated medium. Each measurement was normalized to colony weight. Statistical significance was determined by two-tailed t-tests. * P < 0.05, ** P < 0.01, ns = not significant, ND = not detected.

The pattern of metabolite production on MSgg with and without gluconate was quite different from that of MSN, potentially because MSgg promotes biofilm formation at baseline while MSN does not. Of the five metabolites that increased on MSN gluconate, only surfactin was also significantly increased on MSgg gluconate (Figure 6A). Pulcherrimin, bacillaene, and plipastatin were not significantly different between MSgg conditions (Figure 6B-D), while SKF and subtilosin were not detected on MSgg (Figure 6E-F). Bacillibactin production was significantly lower in the presence of gluconate on both MSN and MSgg (Figure 6G). Of the typical *B. subtilis* specialized metabolites, only bacilysin was not detected in any of our media conditions. The consistent decrease in bacillibactin under both gluconate-containing conditions (Figure 6G) suggested that iron was more readily available in the presence of gluconate, thereby reducing the need for siderophore production. This was also consistent with the fact that pulcherrimin was significantly higher in MSN gluconate and trended higher in MSgg gluconate (Figure 6D): pulcherrimin has been shown to extracellularly chelate excess iron (49).

We next examined whether gluconate itself contributed a substantial amount of iron to the medium by quantifying the iron content of MSN glucose and MSN gluconate using inductively coupled plasma mass spectrometry (ICP-MS), an analytical technique for detecting metals. While baseline iron concentrations in the media with no cells added were higher with gluconate than glucose (Figure 6H), it only increased to 0.17 µM, which is approximately 295X lower than the 50 µM iron added to MSgg. Therefore, the decrease in bacillibactin and observed phenotypic changes, particularly in MSgg, are unlikely to be explained by the small amount of iron contributed by gluconate. We also used ICP-MS to quantify iron levels in whole colony biofilms grown with and without gluconate. On MSN gluconate, iron trended higher but was not significantly different from colonies grown on MSN glucose (Figure 6I). In MSgg, iron was not different with or without gluconate. Overall, these results demonstrate that *B. subtilis* needs less siderophore production to obtain the same amount of iron when it is grown on gluconate, indicating that gluconate may facilitate iron access.

### Gluconate increases iron availability to *B. subtilis* independently from bacillibactin and gluconate uptake

Iron is critical for *B. subtilis* biofilm formation (37), and the decrease in bacillibactin detected by LC-MS/MS (Figure 6G) indicates that gluconate may be connected to changes in iron availability. To better understand the connection between gluconate and iron, we constructed single-deletion mutants in *B. subtilis* iron uptake systems and tested their phenotypes in a high throughput screen. We constructed at least one deletion mutant in every iron uptake system (41) and examined the morphology of each mutant on MSgg and MSgg gluconate (Table 1, Figure S13). Interestingly, most deletion strains had a similar phenotype to wild type on both media conditions (Figure S13). However, strains deleted in genes encoding bacillibactin production (*dhbC, dhbE*) were smaller and less wrinkled on MSgg than wild-type *B. subtilis*; the addition of 0.5% gluconate to MSgg alleviated these phenotypes, leading to increased colony size and wrinkling (Table 1, Figure S13). In addition, strains missing genes involved in bacillibactin uptake (*feuA-C*), were essential for growth on MSgg, but grew, albeit poorly, in the presence of 0.5% gluconate (Table 1, Figure S13). Deletion of the *feu* ATPase, *yusV*, reduced wrinkling and growth in both media conditions (Table 1, Figure S13).

**Table 1.**
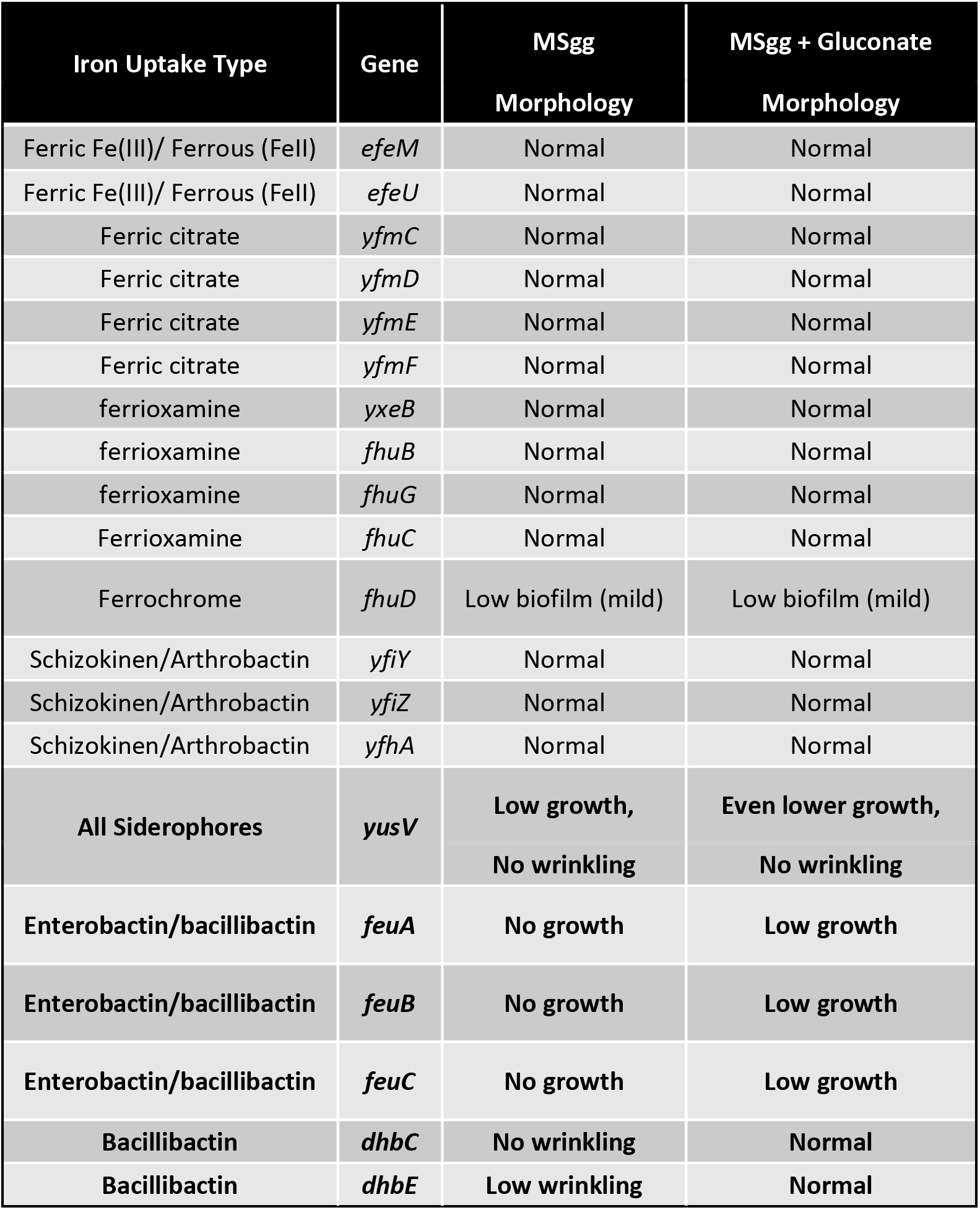
Results from a high-throughput screen of *B. subtilis* 3610 iron uptake deletion strains. . Images of *B. subtilis* 3610 colony biofilms were taken after 48 hours of growth on MSgg with or without 0.5% w/v gluconate. Each deleted gene was replaced with a kanamycin resistance cassette. For each deleted gene, the table lists the type of iron uptake supported by that gene and the colony morphology observed in the screen on both media conditions.

As a follow-up to these high-throughput imaging experiments, we confirmed phenotypes using single-gene deletion strains from each operon identified above (for *dhbA, feuA,* and *yusV*) on both MSN and MSgg agar, which enabled cleaner colony images to be obtained and reduced the possibility of cross-colony interactions (Figure S14). Here we examined the *dhbA* rather than the *dhbC* mutant to eliminate the formation of any bacillibactin intermediates. Results on MSgg were generally consistent with the initial screen, except for *yusV*, which grew better on MSgg gluconate in follow-up experiments, exhibiting the same phenotype as the *feuA* mutant (Figure S14). The *dhbA* bacillibactin mutant showed dramatically more wrinkling and a larger colony diameter on MSgg gluconate (Figure S14). On MSN, *dhbA, feuA,* and *yusV* were all required for growth, with slightly more growth visible on MSN gluconate (Figure S14). The combination of increased growth and wrinkling induced by gluconate, even in the absence of bacillibactin uptake (*feuA* and *yusV* mutants) or when bacillibactin production was eliminated (*dhbA* mutant), indicates that gluconate leads to better iron availability in a manner independent of bacillibactin and its ABC transport system.

Gluconate can form gluconate-iron complexes (61), so we next tested whether its mechanism of providing iron to *B. subtilis* was dependent on the import of iron-bound gluconate. We generated two double-deletion strains of *B. subtilis* that are unable to import gluconate and either cannot make bacillibactin (*gntP*::kan *dhbA*::erm) or cannot make the bacillibactin ABC transporter (*gntP*::erm *feuA*::kan), respectively. When grown on MSgg with and without gluconate, these double mutants had the same phenotype as their respective single deletion *dhbA* and *feuA* strains (Figure 7), indicating that import of gluconate was not required for it to alter iron access.

**Figure 7.**
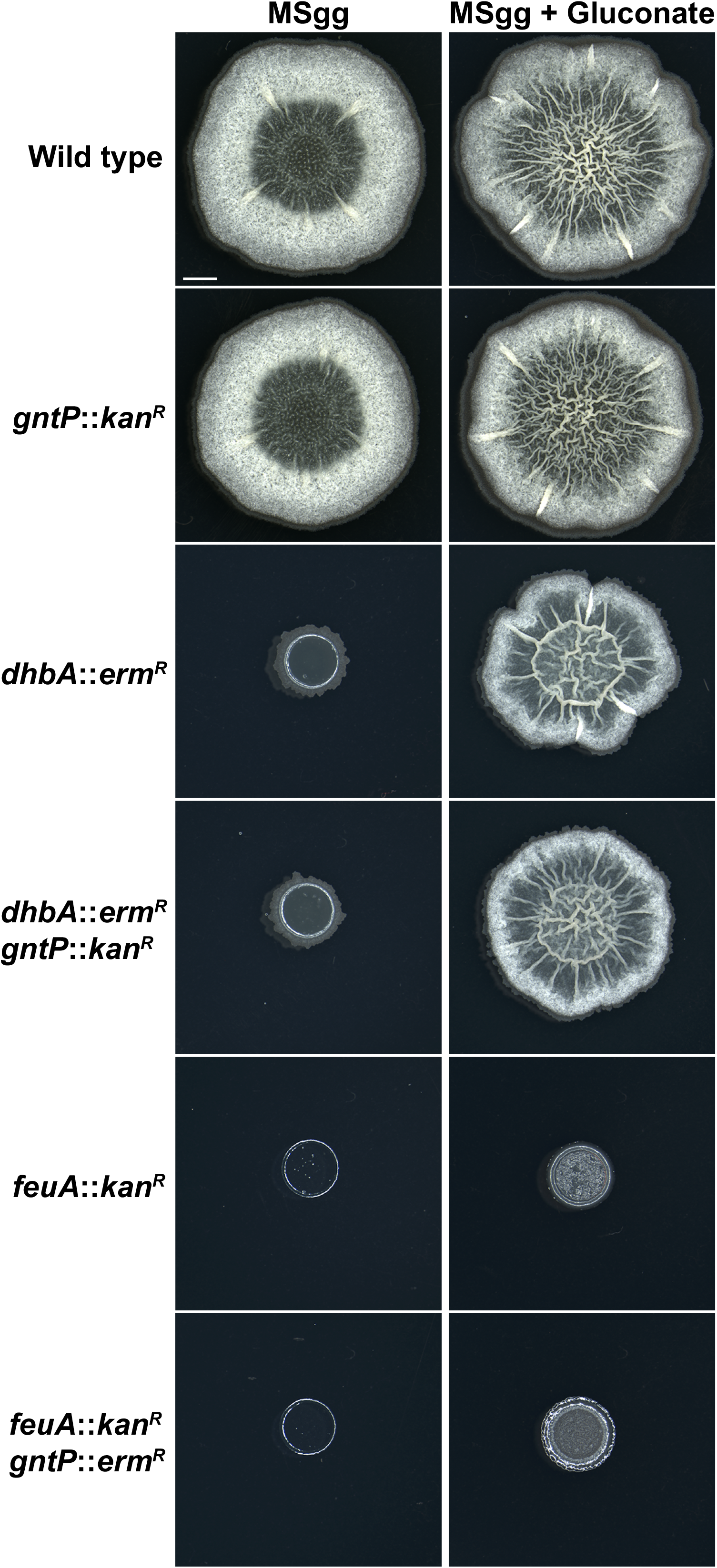
Gluconate uptake is not required to alter iron acquisition in *B. subtilis*. Representative brightfield images (n=2 biological replicates) of wild-type and deletion strains of *B. subtilis* 3610 colony biofilms after 48 hours of incubation on MSgg with or without 0.5% w/v gluconate. Deleted genes are replaced by kanamycin (kan^R^) or erythromycin (erm^R^) antibiotic cassettes. The scale bar indicates 2 mm and is consistent across all images.

Finally, because MSN contains no supplemented iron at baseline, and we suspected that gluconate improves iron access, we tested the responsiveness of *B. subtilis* biofilm colonies on MSN to exogenously added iron. Supplementing ferric iron (FeCl_3_) to the *dhbA*-deficient mutant increased growth at a 1μM concentration and wrinkling at a 100μM concentration on gluconate but not on glucose (Figure S15). Taken together, these results confirmed that gluconate can improve iron availability to *B. subtilis* in a gluconate-import-independent manner, demonstrating a novel interaction between carbon source, biofilm development, and iron accessibility.

## Discussion

Although *B. subtilis* is a well-studied model bacterium, important gaps remain in our understanding of how environmental factors shape its physiology and development (3–5). Here, we used genetic approaches with targeted and untargeted screening and a focus on colony phenotype to uncover novel aspects of *B. subtilis* metabolism, biofilm regulation, and iron acquisition. We have shown that gluconate, a carbon source that is generated by soil microbial metabolism (44–47), induces biofilm gene expression in *B. subtilis* grown in liquid culture (Figure 1) and stimulates wrinkling in colony biofilms (Figure 2). Colony biofilm morphology was assessed on two different media. MSN was selected because it contains no metabolizable carbon and therefore is useful for identifying impacts of gluconate as a sole carbon source. MSgg contains two carbon sources, glycerol and glutamate, and strongly induces biofilm. Strikingly, gluconate enhanced wrinkling in MSgg, showing that it acts on top of an already robust biofilm-inducing condition. Overall, these results show that *B. subtilis* may be sensing gluconate as a signal to indicate either a food source or competitor in its vicinity.

Phenotypic wrinkling in colony biofilms was dependent on genes encoding the canonical biofilm structural components, including TasA and EPS (Figure 3). The hydrophobin BslA was required for wrinkling on MSN, but it was not required on MSgg. Surprisingly, despite the necessity of biofilm components and canonical central biofilm regulators (Figure S6), the expression of genes encoding the matrix components did not increase in response to gluconate when *B. subtilis* was grown as a colony biofilm on agar (Figure 4). It is possible that post-translational regulation is different in the presence of gluconate, or that the metabolic state is very different in the presence of gluconate, despite similar total growth (Figure S4), neither of which would have been captured by the approaches used here. The discrepancy in *tapA* expression between liquid culture and on agar was also quite striking. Much of *Bacillus* biofilm research is conducted either in pellicle (liquid) biofilms or in colony (agar) biofilms (4, 17, 18). These results highlight that conclusions across these two approaches cannot be expected to correlate, especially in studies focused on bacterial metabolism.

The gluconate utilization pathway feeds directly into the pentose phosphate pathway, a central metabolic process that generates ribose-5-phosphate for nucleotide biosynthesis and NADPH for cellular metabolism (53, 62–64). Here, we tested whether metabolism of gluconate was required to induce wrinkling in *B. subtilis* colony biofilms. All genes in the gluconate utilization (*gnt*) operon were required for normal growth on gluconate as a sole carbon source (Figure 5, S8), but they were phenotypically similar to wild type *B. subtilis* on MSgg with and without gluconate (Figure 5). Gluconate is therefore likely to act as an extracellular signal to induce phenotypic wrinkling, a finding that is not unprecedented given that *B. subtilis* biofilm formation is responsive to extracellular specialized metabolite signals (65) and to plant polysaccharides (28, 29). We attempted to identify a signal transduction pathway by testing deletion strains of canonical biofilm (*kin*; Figure S10) genes and extracellular signaling peptides (*rap* and *phr* genes; Figure S11, Figure S12) that are known to be responsive to environmental signals (55, 66–68). However, all of these deletion strains either wrinkled in response to gluconate or had severe biofilm deficiency in the absence of gluconate. One potential explanation for this result is that the *rap* and *phr* genes are redundant, so if gluconate signals via multiple pathways, we would not have detected that here. Additionally, it is possible that either *phrI* or *phrK* are important for gluconate signaling, but that cannot be untangled with the deletion strains used here, because both genes are also essential for biofilm formation in the absence of gluconate.

*B. subtilis* biofilm development is characterized and regulated by the production of specialized metabolites (56, 57, 65). These metabolites act as self-signals that increase in a predictable pattern during biofilm development. To understand how gluconate alters *B. subtilis* biofilm development, we quantified these specialized metabolites by LC/MS-MS (Figure 6). Overall, gluconate altered specialized metabolites differently in MSN and MSgg, and most metabolites did not show the same significance and directional change in both media. Only surfactin, which was increased by gluconate, and bacillibactin, which was decreased by gluconate, were significantly different in both media. The decrease of bacillibactin in the presence of gluconate indicated a decreased need for iron scavenging. Furthermore, pulcherrimin, which can chelate extracellular iron and is used by *B. subtilis* to control excess iron and oxidative stress (51, 52), was significantly increased on MSN gluconate and trended upwards on MSgg gluconate, indicating a potential excess of iron in the presence of gluconate. The secretion of pulcherrimin is also a mechanism to reduction iron access by competitors (51, 52), further supporting the idea that *B. subtilis* may react to gluconate as it would to a competitor. To rule out the possibility that gluconate was adding a significant amount of iron to the media, we used ICP-MS to quantify iron in MSN glucose and MSN gluconate (Figure 6). While gluconate contained more iron than glucose, the amount was 295-fold lower than the iron in MSgg. We also quantified iron in whole colony biofilms and found that gluconate did not increase the amount of colony-associated iron. Together, these results show that *B. subtilis* can acquire the same amount of iron in the presence of gluconate but does not require as much bacillibactin to do so.

Given the apparent increase in iron availability to cells grown on gluconate, we tested a suite of iron uptake mutants, with at least one gene deletion in every known iron uptake system in *B. subtilis* (41), to determine whether any specific iron uptake systems were required for the response to gluconate. While the majority of iron deletion strains wrinkled in response to gluconate (Figure S13), we saw that when iron uptake was limited by elimination of bacillibactin production or uptake, colonies grown on gluconate both grew better (*feu* transporter mutants) and wrinkled more (*dhb* bacillibactin mutants) in the presence of gluconate (Figure S13, Figure S14) (35). These effects were not dependent on the import of gluconate into the cell, as demonstrated by an identical phenotypic response between single gene deletion strains deficient in bacillibactin production or uptake, and double deletion strains that were also deficient in gluconate import (Figure 7). Finally, we demonstrated that additional iron in the media overcame the loss of bacillibactin when gluconate but not glucose was the sole carbon source, leading to increased growth and wrinkling (Figure S15). We have demonstrated here that the presence of gluconate in the environment allows for additional iron uptake, suggesting that gluconate facilitates bacillibactin-independent iron uptake without the need for direct uptake of gluconate. Indeed, the biofilm matrix itself is known to bind iron and make it accessible to cells, and it is possible that gluconate is involved with or enhances this phenomenon (38).

The relationship between biofilm and iron in *B. subtilis* is complex. Iron is required for robust biofilm formation and maturation, and the biofilm matrix acts as a reservoir for iron (37–39). However, there is no evidence that iron itself is a signal to induce biofilm formation in *B. subtills*, although uptake of pirated siderophores promotes sporulation, a process that is regulated by the same central regulator (Spo0A) as biofilm formation (40). It is therefore unlikely that iron accessibility is the sole mechanism by which gluconate increases biofilm in *B. subtilis*, but iron access is one important component of the response to gluconate.

Overall, we have shown that gluconate induces biofilm wrinkling in *B. subtilis*, that this wrinkling is independent of gluconate utilization, and that gluconate facilitates environmental iron uptake. These multifaceted results show that the response to gluconate is complex and affects many aspects of *B. subtilis* biology. Future work will focus on identifying the signal that induces biofilm in response to gluconate and better understanding of the mechanism(s) by which gluconate increases iron uptake.

## Materials & Methods

### Bacterial Strains & Media

All bacterial strains in this manuscript are listed in Table S3. *B. subtilis* was routinely cultured on lysogeny broth (LB) agar (10g/L tryptone, 5 g/L yeast extract, 5 g/L NaCl, 1.5% agar), minimal salts nitrogen (MSN): 5 mM potassium phosphate buffer (pH 7), 0.1 M morpholinepropanesulfonic acid (MOPS, pH 7), 2 mM MgCl_2_, 50 µM MnCl_2_, 1 µM ZnCl_2_, 2 µM thiamine, 700 µM CaCl_2_, 0.2% NH_4_Cl, and Minimal Salts Glycerol Glutamate (MSgg): 5 mM potassium phosphate buffer (pH 7), 0.1 M morpholinepropanesulfonic acid (MOPS, pH 7), 50 µM FeCl_3_, 2 mM MgCl_2_, 50 µM MnCl_2_, 1 µM ZnCl_2_, 2 µM thiamine, 700 µM CaCl_2_, 20% v/w glycerol, 20% w/v glutamate. MSN and MSgg were supplemented with the indicated carbon source in each experiment. 1.5% agar was added for colony biofilm experiments, and 30 mL per plate was dispensed. Plates were opened and dried for 10 minutes in a biosafety cabinet prior to inoculation.

### Bacterial Culture

*B. subtilis* was routinely struck onto LB agar and grown at 30°C for 16 hours prior to inoculation of liquid cultures or colony biofilms. Cells were then scraped and washed twice (2 minutes, 10,000 rpm centrifuge steps) in MSN with no added carbon source. For liquid assays in 96-well plates, cells were then diluted to a final OD_600_ of 0.05. For colony biofilm experiments on agar, cells were diluted to a final OD_600_ of 0.5, and 2 µL of this suspension was dropped onto the center of an MSN or MSgg agar plate containing the added carbon source(s) of interest. Plates were dried open on the bench for 5-10 minutes and then placed at 30°C. Incubator humidity levels were maintained at 50-60%.

### Carbon Source Screen

Phenotype microarray carbon utilization plates PM1 and PM2A were purchased from Biolog (Hayward, CA). 120 µL MSN without an additional carbon source was added to each well of these plates and allowed to equilibrate at room temperature for 1 hour. Carbon sources were then homogenized with a pipette tip and 100uL was transferred to a Nunclon Delta Surface 96-well plate. The luminescent *tapA* reporter strain *B. subtilis* 3610 *sacA*::P*_tapA_*-LUXABDCE was grown and washed as described above and was added at a final OD_600_ of 0.05 to each well. Plates were placed in a 30°C incubator and double-orbital shaken at 280 rpm. Luminescence was measured at inoculation and after 8, 12 and 24 hours of culture. The OD_600_ and luminescence values detected at inoculation were background subtracted from detected values at each subsequent time point. To confirm initial carbon source hits, a dehydrated form of each carbon source was purchased, dissolved at 0.5% v/w in MSN, and sterile filtered. *B. subtilis* 3610 *sacA*::P*_tapA_*-LUXABDCE was prepared for each carbon source, as described above in Bacterial Culture. Cells were then diluted to a final OD_600_ of 0.05 in each carbon source of interest and 200 µL was inoculated into each well of a white Greiner Bio-one 96-well µClear plate. Luminescence and OD_600_ measurements were taken hourly on a BioTek Synergy H1 instrument in top-read mode with full-spectrum emission detection. Luminescence was acquired with a 1 s integration time, extended dynamic range enabled, a 4.5 mm read height, and gain of 135. Plates were double-orbital shaken at 280 rpm at 30°C.

### CFU and Sporulation Assays

*B. subtilis* was inoculated in 200 µL liquid culture in white Greiner Bio-one 96-well µClear plates at an OD_600_ of 0.05 and double-orbital shaken at 280 rpm at 30°C. At each timepoint, three wells were combined into a 1.5 mL Eppendorf tube. Cell clumping was disrupted with a QSonica sonicator for 12 s total; 1 s pulse followed by 1 s pause at an amplitude of 15%. For sporulation assays, 100uL was removed from each sample and heated in a thermocycler at 80°C for 20 minutes to kill vegetative cells. Heat treated and non-heat-treated cells were serially diluted in PBS, plated on LB and incubated at 30°C overnight. Colony forming units (CFUs) were counted and the percentage of spores was calculated as ([CFU spores]/[CFU total population])*100. For CFU quantification in colony biofilms, whole colonies were scraped into 1mL of phosphate buffered saline (PBS) from the indicated carbon source and time point. Colonies were then needle-sheared with a 23G needle and 1mL syringe, followed by sonication as described above. Cells were serially diluted and plated on LB agar for CFU quantification.

### Imaging

Photos were taken with a Nikon D7000 digital camera with an AF MICRO NIKKOR 60 mm 1:2.8 D lens unless noted otherwise. Microscopy images were taken on a Leica M165 FC dissecting microscope. Brightfield image exposure was 100 ms. Fluorescence images were taken with a YPET ET filter with a EL6000 light box and 5 s exposure. For fluorescent image quantification, a paired brightfield and fluorescent image was taken of each colony. Fluorescent reporter intensity was quantified in ImageJ. First, a mask was generated from the corresponding brightfield image using the ImageJ wand tool. The mask was then applied to the paired fluorescent image, and mean fluorescence intensity was quantified. Colony images were adjusted for figure display in Photoshop and figures were assembled in Adobe Illustrator. The brightness and contrast of colony images was linearly adjusted. Image size was decreased and images were cropped. Image scale bar size was calculated from the Leica metadata using the appropriate size adjustment applied to each image.

### Mutant Strain Construction in *B. subtilis*

All new gene deletions in this manuscript were constructed using phage transduction, as previously described (69). Donor strains harboring genes replaced by an antibiotic cassette were obtained from the Bacillus Genetic Stock Center in the *B. subtilis* 168 strain background. To prepare phage, a donor strain was struck overnight on LB agar. The strain was then resuspended in 3 mL TY media (LB supplemented with 10mM MgSO_4_ and 100 M MnSO_4_). Cells were grown at 37°C on a roller until late-log phase and then infected with SPP1 phage for 15 minutes at 37°C. Cells were suspended in 3mL 0.5% TY top agar and top agar was spread evenly onto an LB plate, which was incubated overnight at 37°C. Top agar was collected into 5mL TY broth the next day and vortexed to homogenize. Samples were centrifuged at speed 5 in a clinical centrifuge for 7 minutes, and the supernatant was transferred to a new tube. MgSO_4_ was added to 10mM and DNase I to 25 μg/mL. Samples were incubated for 5 minutes at room temperature and then filtered through a 45 μM syringe filter. Lysate was stored at 4°C. To infect a recipient strain with phage, the recipient was grown overnight on an agar plate at 30°C and then subcultured in TY broth at 37°C until early stationary phase. Diluted phage lysate was added to 1mL of cells and incubated at 37°C for 30 minutes. Infected cells were plated on LB supplemented with 10mM citrate and an antibiotic corresponding to the appropriate antibiotic cassette (1 µg ml^−1^ erythromycin or 25 µg ml^−1^ lincomycin and kanamycin 5 µg ml^−1^). Plates were incubated at 37°C overnight, and single colonies were substruck twice on LB plus antibiotic to remove phage.

### Flow Cytometry

Whole colonies or pellicles were fixed as previously described (70). Briefly, cells were collected into 1 mL PBS in a 1.5 mL Eppendorf tube. Cells were then sheared with a 23G needle and pelleted at 13,000 rpm for 2 minutes. Supernatant was removed and cells were resuspended in 200 µL of 4% paraformaldehyde. Cells were incubated at room temperature for 7 minutes. Cells were centrifuged as above, paraformaldehyde was removed, and cells were washed with 1 mL PBS. Cells were centrifuged again, PBS was removed, and cells were resuspended in GTE buffer (47 mL PBS, 2.5 mL 20% glucose, 0.5mL 0.5 M EDTA). All steps were performed at 4°C unless otherwise indicated, and cells were stored at 4°C after fixation. Directly prior to flow cytometry, cells were sonicated twice as described above, with a minimum 30 s rest on ice between sonication steps. Flow cytometry was performed on a BD LSRFortessa with BD FACSDiva v9.0 software. A total of 100,000 events were collected per sample. The instrument was configured with 405-nm violet, 445-nm blue-violet, 488-nm blue, and 561-nm yellow-green excitation lasers. Forward scatter (FSC), side scatter (SSC), and FITC fluorescence were collected. FITC fluorescence was excited by the 488-nm blue laser and detected using a 530/30 bandpass filter. Photomultiplier tube voltages were set to 500 V (FSC), 350 V (SSC), and 550 V (FITC) with acquisition thresholds of 400 FSC and 500 for SSC. Analysis of flow cytometry data was performed in FlowJo, with a wild-type non-fluorescent *B. subtills* control used to determine the cutoff between fluorescent and non-fluorescent populations.

### ONPG assay

Gene expression in the EPS reporter strain *B. subtilis* 3610 *amyE::*P*epsA-lacZ* was quantified by *o*-nitrophenyl-β-D-galactopyranoside (ONPG) assay as previously described (28). Briefly, whole colonies were resuspended in 1mL PBS and sonicated as described above. The OD_600_ of the sample was measured, and samples were then resuspended in Z buffer (40 mM NaH_2_PO_4_, 60 mM Na_2_HPO_4_ 1 mM MgSO_4_, 10 mM KCl, 38 mM β-mercaptoethanol, 200 µg/mL Lysozyme) at 30°C for 15 minutes. 200 uL of ONPG was added and samples were vortexed to homogenize. Once reactions turned yellow, 500 µL of 1M Na_2_CO_3_ was added to each sample to stop the reaction. Samples were centrifuged at 1300 rpm for 1 minute, and the OD_420_ of the soluble fraction was measured. B-galactosidase activity was calculated as (OD_420_/(reaction time (min) * OD_600_))*1000. ONPG values from a paired wild type strain without the *lacZ* reporter were subtracted from the final quantification.

### ICP-MS

Elemental analysis was performed by inductively coupled plasma mass spectrometry (ICP-MS) at the UMass Amherst ICP-MS Core Facility. Fresh liquid MSN with 0.5% glucose or gluconate was used to quantify baseline iron in the media. To quantify colony biofilm associated iron, whole colonies of wild type *B. subtilis* were scraped from agar after 48h growth at 30°C, weighed, and frozen. Iron quantities in parts per million (ppm) were normalized to colony weight.

### Metabolomics spectrometry analysis

Colony biofilms and agar were collected after 48 hours growth at 30°C by coring with a size-9 cork borer (McMaster-Carr). For each biological replicate, 4 colonies were collected into a total volume of 16 mL 80% methanol in an Erlenmeyer flask. Samples were shaken at 150 rpm at room temperature for 2 hours. The liquid part of the sample was aspirated with a syringe and needle and filtered over a 0.22 µM PVDF syringe filter to sterilize. The whole sample was transferred to a 20 mL glass vial and dried over a speed vacuum. Liquid chromatography-tandem mass spectrometry (LC-MS/MS) data were acquired using a Q-Exactive HF-X quadrupole-Orbitrap mass spectrometer (ThermoFisher Scientific) with a heated electrospray ionization (HESI) source coupled to a Vanquish ultrahigh-performance LC (UPLC) system (ThermoFisher Scientific). Samples were reconstituted in 10% methanol for analysis. Injections of 5 μl were performed on an Acquity UPLC BEH C_18_ column (1.7 μm, 2.1 by 50 mm; Waters) with a flow rate of 0.5 mL/min using the following binary solvent gradient of A (H_2_O, 0.1% formic acid added) and B (CH_3_CN, 0.1% formic acid added): initial isocratic composition of 90:10 (H_2_O-CH_3_CN) for 1.0 min, followed by a series of linear decreases in A: to 60% A from 1 to 1.5 min, to 60% A from 1.5 to 2.5 min, to 10% A from 2.5 to 6.5 min, and to 0% A from 6.5 to 9.5 min. The level of A was increased from 0 to 90% from 9.5 to 10.5 min, and held at 90% A from 10.5 to 13 min. The first minute of analysis was diverted to waste. The positive ionization mode was utilized over a full scan of *m/z* 200 to 2,000 with the following settings: capillary temperature, 270°C; tube lens offset, 45 V; spray voltage, 3.80 kV; sheath gas flow and auxiliary gas flow, 53 and 14 U, respectively. Sweep gas was set to 3 U. Resolution was set to 60,000 with an automatic gain control target of 1E6. The top 5 most intense ions in each scan were fragmented at stepped normalized collision energies of 15, 35, and 60 V. Ions were dynamically excluded for 5 sec after initial fragmentation. MS/MS spectra were collected for each sample using data-dependent acquisition, with resolution set to 30,000 and an automatic gain control target of 1E5. Extracted ion chromatograms were obtained from the XCalibur software (ThermoFisher Scientific).

### Metabolomics data processing

Raw data files were analyzed, aligned, and filtered using MZmine 4.4.3 software (71, 72) (http://mzmine.github.io/). All parameters used for feature finding are listed in Table S4. Briefly, MS1 and MS2 masses were detected in each file and chromatograms were made for each feature. Chromatograms were smoothed and deconvoluted, and isotopic peaks were grouped. The join algorithm was used to integrate all the chromatograms into a single data matrix using the following parameters: the balance between *m/z* and retention time was set at 10.0 each, *m/z* tolerance was set at 0.005 or 10 ppm, and retention time tolerance was defined as 0.30 min. The resulting data matrix was gap-filled, filtered to ensure each feature was detected in at least 5 files, and all features with an associated MS2 spectrum were exported for analyses.

### High-throughput screen of iron deletion strains

A high throughput morphology screen of *B. subtilis* 3610 iron uptake deletion strains was performed using a Singer ROTOR HDA instrument for high throughput replication in a 96-well format. Deletion strains were constructed as described above. Strains were inoculated into 100 µL LB broth in a 96-well Nunclon Delta Surface 96-well plate and cultured at 30°C with double orbital shaking at 280 rpm for 24 hours. 100 µL of LB + 15% glycerol was then added to each well, and plates were sealed and stored frozen at -80°C. Frozen culture was inoculated into 200 µL LB in a Nunclon Delta Surface 96-well plate using a 96-well replicator and cultured for 24 hours as above. MSgg agar plates with and without 0.5% gluconate were prepared in Singer PlusPlates with 55 mL media per plate. Plates were dried for 10 minutes with the lid off in a biosafety cabinet prior to inoculation. A Singer ROTOR instrument was used to replicate cultured LB plates to MSgg and MSgg plus 0.5% w/v Gluconate. Plates were dried on the bench with lids removed for 5-10 minutes and then incubated at 30°C for 48 hours. Images were acquired with a Canon EOS R100 digital camera fitted with a Canon EF-S 60 mm f/2.8 Macro USM lens.

## Author Contributions

CE Price: Conceptualization, Data curation, Formal Analysis, Investigation, Methodology, Validation, Visualization, Writing – original draft, Writing – review & editing

A Sadlon: Data curation, Investigation, Methodology, Validation

A Bradley: Data curation, Formal Analysis, Investigation, Methodology, Validation

C Perez: Investigation, Methodology

EA Shank: Conceptualization, Funding acquisition, Project administration, Resources, Supervision, Visualization, Writing – review & editing

## Acknowledgements

We are grateful to Amir Mitchell for use of his lab’s instrumentation for high throughput experiments. This research was supported by funds provided by the National Institutes of Health (R35GM145261 to EA Shank),

